# VaxArray Immunoassay for the Multiplexed Quantification of Poliovirus D-Antigen

**DOI:** 10.1101/2021.08.30.458257

**Authors:** Erica D. Dawson, Amber W. Taylor, James E. Johnson, Tianjing Hu, Caitlin McCormick, Keely N. Thomas, Rachel Y. Gao, Rahnuma Wahid, Kutub Mahmood, Kathy L. Rowlen

## Abstract

Next generation poliovirus vaccines are critical to reaching global poliovirus eradication goals. Recent efforts have focused on creating inactivated vaccines using attenuated Sabin strains that maintain patient safety benefits and immunogenicity of conventional inactivated vaccines while increasing manufacturing safety and lowering production costs, and on developing novel oral vaccines using modified Sabin strains that provide critical mucosal immunity but are further attenuated to minimize risk of reversion to neurovirulence. In addition, there is a push to improve the analytical tools for poliovirus vaccine characterization. Conventional and Sabin inactivated poliovirus vaccines typically rely on standard plate-based ELISA as *in vitro* D-antigen potency assays in combination with WHO international standards as calibrants. While widely utilized, the current D-antigen ELISA assays have a long time to result (up to 72 hours), can suffer from lab-to-lab inconsistency due to non-standardized protocols and reagents, and are inherently singleplex. For D-antigen quantitation, we have developed the VaxArray Polio Assay Kit, a multiplexed, microarray-based immunoassay that uses poliovirus-specific human monoclonal antibodies currently under consideration as standardized reagents for characterizing inactivated Sabin and Salk vaccines. The VaxArray assay can simultaneously quantify all 3 poliovirus serotypes with a time to result of less than 3 hours. Here we demonstrate that the assay has limits of quantification suitable for both bioprocess samples and final vaccines, excellent reproducibility and precision, and improved accuracy over an analogous plate-based ELISA. The assay is suitable for adjuvanted combination vaccines, as common vaccine additives and crude matrices do not interfere with quantification, and is intended as a high throughput, standardized quantitation tool to aid inactivated poliovirus vaccine manufacturers in streamlining vaccine development and manufacturing, aiding the global polio eradication effort.

## 1. INTRODUCTION

Effective vaccines against poliomyelitis have been available since 1955, when the original Salk inactivated polio vaccine received licensure in the United States.^1^ Both inactivated polio vaccines (IPV) and live attenuated oral polio vaccines (OPV) have been the cornerstone of global polio vaccination initiatives since their original introductions in the late 1950’s and early 1960’s. Consistent global vaccination efforts and a focus on polio eradication by the World Health Organization (WHO) Global Polio Eradication Initiative (GPEI) starting in 1988 have successfully led to certification of global eradication of both wild poliovirus type 2 in 2015 and type 3 in 2019.^2,3^

Given the ability of the attenuated Sabin strains utilized traditionally in the manufacturing of OPV to cause outbreaks in regions of low population immunity by vaccine-derived polioviruses (VDPVs), there are continued discussions and efforts to minimize and eliminate the use of OPV in the post-eradication era.^4,5^ In fact, following the eradication of poliovirus type 2, a coordinated worldwide ‘switch’ from the trivalent OPV containing types 1, 2, and 3 to a bivalent OPV containing only types 1 and 3 was successfully executed in April 2016 to minimize the risk of reintroduction of live type 2.^2,5^ On the other hand, while conventional IPV (cIPV) manufactured from wild-type viruses are inactivated and therefore do not have the ability to replicate post-vaccination, cIPVs must be manufactured under high biosafety conditions and carry the risk of outbreaks due to unintentional mishandling or breach of containment.^5–7^

Inactivated vaccines based on the attenuated Sabin strains, dubbed Sabin IPV or sIPV, have been the focus of significant efforts since the 1980’s, with some countries now having licensed sIPV vaccines, including Japan’s use of trivalent sIPV in a combination vaccine since 2012 and China’s use of standalone trivalent sIPV starting after approval in 2015.^5,8^ In December 2020, the LG Chem Eupolio Inj. vaccine was the first sIPV vaccine to achieve WHO prequalification.^9^ The use of attenuated strains reduces the biosafety risk of accidental exposure during manufacturing, they are more suitable for manufacturing in in lower- and middle-income countries, and literature indicate that cost reductions over cIPV may be realized.^8,10^ In addition, because sIPV are inactivated, the risk of seeding VDPVs is eliminated.

The advent of sIPV requires a more reliable *in vitro* potency assay for assessing D-antigen content in these vaccines. The typical *in vitro* potency assay for conventional IPV is a D-antigen ELISA using appropriate WHO standards, and while efforts have been made to standardize the recommended protocol, historically vaccine manufacturers have developed their own in-house ELISAs with wide variability in reagent composition, protocol, and performance.^11^ Recently PATH funded the development of human monoclonal antibodies (mAbs) for each poliovirus serotype, with the goal being standardized reagents equally suitable for both sIPV and cIPV D-antigen ELISAs. In work published by Kouiavskaia et al., these PATH-funded mAbs with serotype-specificity were used to capture poliovirus D-antigen and a “universal” pan-poliovirus D-antigen mAb was used for detection in a standard plate-based ELISA.^12^ The 3-day assay exhibited robust analytical performance and the capability of accurately measuring potency of both cIPV and sIPV materials using appropriate standards.

In this work, we present the integration of these recently-developed human monoclonal antibodies against poliovirus D-antigen into a microarray immunoassay platform to achieve multiplexing, increased standardization, and a significantly faster time to result than ELISA methods. The analytical performance metrics for sIPV and cIPV are presented, including sensitivity, specificity, accuracy, precision, absence of cross-reactivity from common vaccine additives, along with accuracy of the method relative to an analogous 3-day plate-based ELISA.

The ≤ 3 hour VaxArray Polio Assay exhibits nearly equivalent performance to ELISA but enables simultaneous analysis of all serotypes in multivalent formulations, including combination vaccines, using a standardized off the shelf kit.

## 2. MATERIALS AND METHODS

### 2.1 VaxArray Poliovirus Assay Design

Three human monoclonal antibodies, specific to poliovirus D-antigen serotypes 1, 2, and 3, were printed on the array at optimized concentrations, and the universal human monoclonal detection antibody was conjugated to a fluorescent dye utilizing a commercially available conjugation kit with a VaxArray-compatible fluorophore (~530 nm excitation, ~630 nm emission).

A schematic of the VaxArray polio assay microarray and detection principle is shown in **Figure 1**.

**Figure 1.**
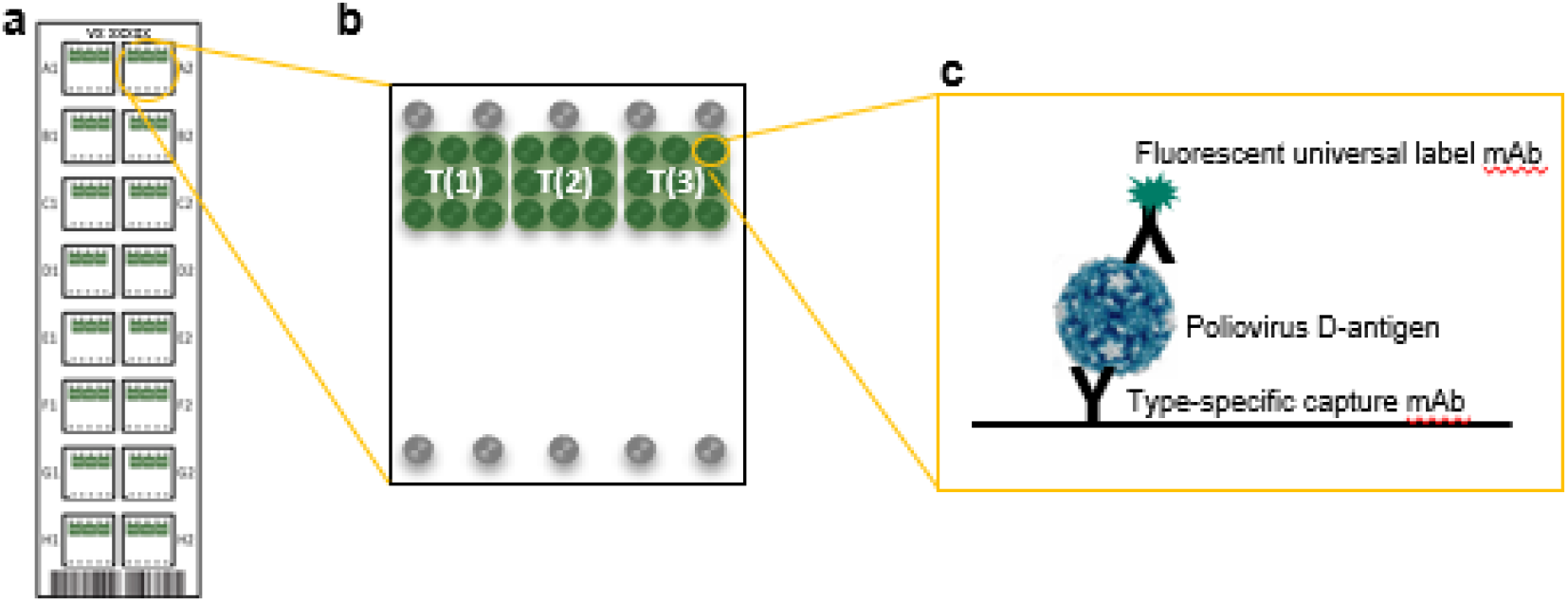
(a) Schematic representation of the VaxArray Polio Assay Kit microarray slide showing 16 replicate microarrays, (b) individual microarray layout showing 9 replicate spots in green for each serotype T(1), T(2), and T(3), and fiducial markers in grey, (c) general assay detection scheme showing serotype-specific capture with a monoclonal antibody and universal (pan-serotype) detection labeling.

In **Figure 1a**, the 16 replicate arrays included on the microarray slide are shown. Each microarray, as shown in **Figure 1b**, contains 9 replicate spots of each of the three serotype-specific monoclonal antibody captures T(1), T(2), and T(3), as well as fiducial marker spots shown in gray for array location by the analysis software. The general assay detection principle is shown in **Figure 1c** in which the serotype-specific monoclonal capture antibodies are immobilized on the functionalized microarray glass surface. Poliovirus D-antigen is then captured, excess antigen washed away, and captured antigen is labeled with a fluorescent monoclonal antibody universal for all three poliovirus D-antigen serotypes.

### 2.2 VaxArray Poliovirus Assay Procedure

The VaxArray Polio Assay Kit (VXPL-9000, InDevR, Inc.) contains two microarray slides, with 16 replicate microarrays per slide, an optimized Protein Blocking Buffer (PBB), and Wash Buffer concentrates. Prior to use, microarray slides were equilibrated to room temperature for 30 min in the provided foil pouch. Standards and samples were diluted in PBB, applied to the microarray, and allowed to incubate in a humidity chamber (VX-6203, InDevR, Inc.) on an orbital shaker (SCI-O180-S, Scilogex) at 80 rpm for 2 hr.

During sample incubation, VaxArray Poliovirus Detection Label (VXPL-7660, InDevR, Inc.) was diluted to 1x in PBB. Samples were removed, and detection label added to the microarray and incubated on an orbital shaker at 80 rpm for 30 min in a humidity chamber. Following label incubation, sequential washes in Wash Buffer 1, Wash Buffer 2, 70% Ethanol (BP82031GAL, Fisher Scientific), and finally ultrapure water (Option Q15, ELGA) were performed. Slides were dried using the VaxArray Slide Drying Station (VX-6208, InDevR, Inc) and imaged using the VaxArray Imaging System (VX-6000, InDevR, Inc.). Any deviations from this standard protocol are noted in the subsequent sections.

### 2.3 Poliovirus Materials and Sample Preparation

The sIPV WHO international standard (17/160) and cIPV WHO international standard (12/104) were obtained from NIBSC. Oral Poliovirus (OPV) WHO international standards were obtained from NIBSC (16/196, 15/296, and 16/202). Purified monovalent bulk sIPV materials and research quantities of the antibodies used on the microarray were obtained from PATH and were calibrated against the sIPV WHO international standard to assign known concentrations. Vaccines including IPOL (NDC 49281-860-78, Sanofi Pasteur), Pentacel (NDC 49281-510-05, Sanofi Pasteur), Daptacel (NDC 49281-286-10, Sanofi Pasteur), EngerixB (NDC 58160-821-11, Sanofi Pasteur), and Pediarix (NDC 58160-811-52, GSK) were purchased from Global Sourcing Initiative (Miami, FL). Materials for interference testing included 2-phenoxyethanol (2-PE) (77699-250ML, Sigma-Aldrich, St Louis, MO), sodium citrate tribasic dihydroxide (C8532-100G, Sigma-Aldrich), and Imject Alum (77161,Thermo Fisher Scientific, Waltham, MA) and Vero cells. Vero cell material was obtained from 0.25% Trypsin-EDTA (25200056, Thermo-Fisher) harvested Vero cell (CCL-81, ATCC, Manassas, VA) culture grown in Medium-199 (11150059, Gibco, Waltham, MA) supplemented with 5% fetal bovine serum (A3160401, Gibco), 2 mM L-Glutamine (A2916801, Gibco), and 1X Penicillin-Streptomycin (15140148, Gibco). All samples for VaxArray Polio Assay analysis were diluted to final testing concentrations in PBB unless otherwise noted.

### 2.4 Specificity

#### Antibody Specificity for Poliovirus Types 1, 2, and 3

Serotype specificity of the capture antibodies was verified with monovalent sIPV analyzed at high concentrations of 80/40/80 D-antigen (D-Ag) units/mL for types 1/2/3 to ensure that any low-level cross-reactivity between serotypes would be observed. Detection label only was also run as a blank to ensure no direct binding of the detection label to the captures.

#### D-Antigen Specificity

The trivalent sIPV WHO standard was prepared at concentrations of 2.5 D-Ag units/mL in each serotype. An aliquot was left untreated, and another series of aliquots were heat treated at 56°C for varying amounts of time to allow conversion of the immunogenic D-antigen form to the non-immunogenic C-antigen form. After heat treatment, all the samples were analyzed to assess VaxArray Polio Assay signal as a function of time at 56°C as compared to the untreated sample.

### 2.5 Linear Dynamic Range and Limits of Quantification

For sIPV, lower and upper limits of quantification (LLOQ and ULOQ) were determined in both monovalent and trivalent samples. For cIPV, the LLOQ was determined using IPOL vaccine (Sanofi Pasteur) that was calibrated against the cIPV WHO international standard, and ULOQ was established using the trivalent cIPV WHO international standard, as IPOL contains concentrations too low to probe ULOQ. To determine the LLOQ, 3 samples were prepared at concentrations producing signal slightly above background. To determine the ULOQ, three samples were prepared at concentrations near the upper edge of the linear range based on a 16-point dilution series. Eight (8) replicates of the sIPV samples and 4 replicates of the cIPV samples were run alongside a standard curve of the same material. LLOQ and ULOQ was defined for each serotype and material as the lowest or highest concentration, respectively, at which %RSD of replicates was < 20% and accuracy of replicates was within ± 25% of the expected value. Dynamic range of the assay was expressed as ULOQ/LLOQ.

### 2.7 Precision and Accuracy

To characterize precision and accuracy, a study was conducted in which 3 users analyzed 8 replicates of trivalent sIPV over each of 3 days (3 users x 8 replicates x 3 days = 72 replicates). On each day of testing, purified sIPV monovalent bulk were mixed to create 8 replicate aliquots of a trivalent sample at 5 D-Ag units/mL of each serotype. In addition, a serial dilution of the same trivalent sample was prepared and analyzed by each user on each day as a standard curve alongside the replicates to enable accuracy calculations. Precision was quantified for each user as well as combined over all 3 users and expressed as the %RSD of replicate measurements. To investigate assay accuracy, the 3 standard curves generated by each user were averaged, and then utilized to back-calculate the measured concentrations. Accuracy was calculated as the % of expected concentration (measured value divided by expected value, expressed as a percentage), and again quantified for each user as well as combined over all 3 users.

A single user reproducibility and accuracy study was also conducted for cIPV using IPOL vaccine. The concentrations in the IPOL vaccine (85.9/21.9/77.45 D-Ag/mL for types 1/2/3) were determined by quantification against the WHO cIPV international standard (12/104, NIBSC). Eight (8) replicates of IPOL were run at a 1:17.2 dilution, and this was repeated on three separate days, generating n=24 replicates over the three days.

Lastly, the inter-assay reproducibility and accuracy of quantifying sIPV and cIPV were investigated at low, medium, and high concentrations to compare metrics in different ranges of the response curve. Testing was completed by a single user. For sIPV, 8 replicates for each of three (3) contrived trivalent samples prepared at a low, medium, or high concentration (1.5, 5.0, and 10.0 D-Ag units/mL, respectively) were analyzed. Three dilutions of the trivalent IPOL vaccine (low = 1.5/0.38/1.35, medium = 5/1.27/4.51, and high = 10/2.55/9.02 D-Ag units/mL for types 1/2/3) were prepared to evaluate cIPV. Standard curves of the same material were included in each assay setup to allow quantification.

### 2.8 Standard Plate-Based ELISA

The performance of the VaxArray Polio Assay was compared to the standard plate-based ELISA of Kouiavskaia et al. that utilizes the same human antibodies.^12^ Antibodies were prepared 1:500 in carbonate-bicarbonate coating buffer (SRE0034, Sigma-Aldrich), added to each well of a Costar 96-well Assay plate (3369, Corning), and incubated overnight at +4°C in a humidity chamber. Plates were washed 3x using 0.5% Tween20 in PBS. 100 uL of blocking buffer (3% BSA in PBS) was added to each well and incubated at +25°C for 1-hr. After incubation, plates were again washed, 50 μL of standards and samples in dilution buffer (1% BSA in PBS) added to the appropriate wells, and incubated overnight at +4°C. The next day, the plates were washed as described above. The detection antibody, biotinylated with 7:1 biotin-to-antibody using EZ-Link Sulfo-NHS-LC-Biotinylation Kit (21435, Thermo Fisher Scientific), was prepared in dilution buffer, and 50 μL added to each well. Plates were incubated for 90 min in a +25°C incubator. After incubation, plates were washed, and 50 μL of ExtrAvidin Peroxidase (E2886, Sigma-Aldrich) diluted 1:1000 in dilution buffer was added to each well and incubated for 40 min at +25°C. Plates were washed, 100 μL of TMB (5120-0075, SeraCare, Milford, MA) was added to each well, and incubated on an orbital shaker (80 rpm) in the dark at +25°C for 15 min. TMB STOP-reagent (S5814, Sigma-Aldrich) was added, and plates read at 450 nm on a FLUOstar OPTIMA (BMG LABTECH, Ortenberg Germany) microplate reader.

To compare the VaxArray Polio Assay to the standard plate-based ELISA, a trivalent 2x mixture of sIPV material was prepared in PBS and diluted 1:1 in dilution buffer (for ELISA) or in PBB (for VaxArray). Six additional dilutions were prepared with each sample run in triplicate. The sIPV samples were run alongside the sIPV WHO International Standard for sIPV for quantification. For ELISA, three separate 6-point standard curves were made (ranging from 0.03-1, 0.02-0.625, 0.04-1.25 D-Ag units/mL for types 1, 2, and 3, respectively). For the VaxArray Polio Assay, one single 7-pt curve was prepared over the range from 0.06-20 D-Ag units/mL. The experiment for both assays was repeated twice (total of n=6 replicates for each sample).

### 2.9 Interference

#### Cross-Reactivity in the Absence of Poliovirus

To evaluate cross-reactivity of common vaccine components, Daptacel (containing DTaP), the *H. influenza* component of Pentacel, and ENGERIX-B (Hepatitis B) were tested. An excerpted list of the critical ingredients in these vaccines is provided in **Table 1**.^13–17^

**Table 1:**
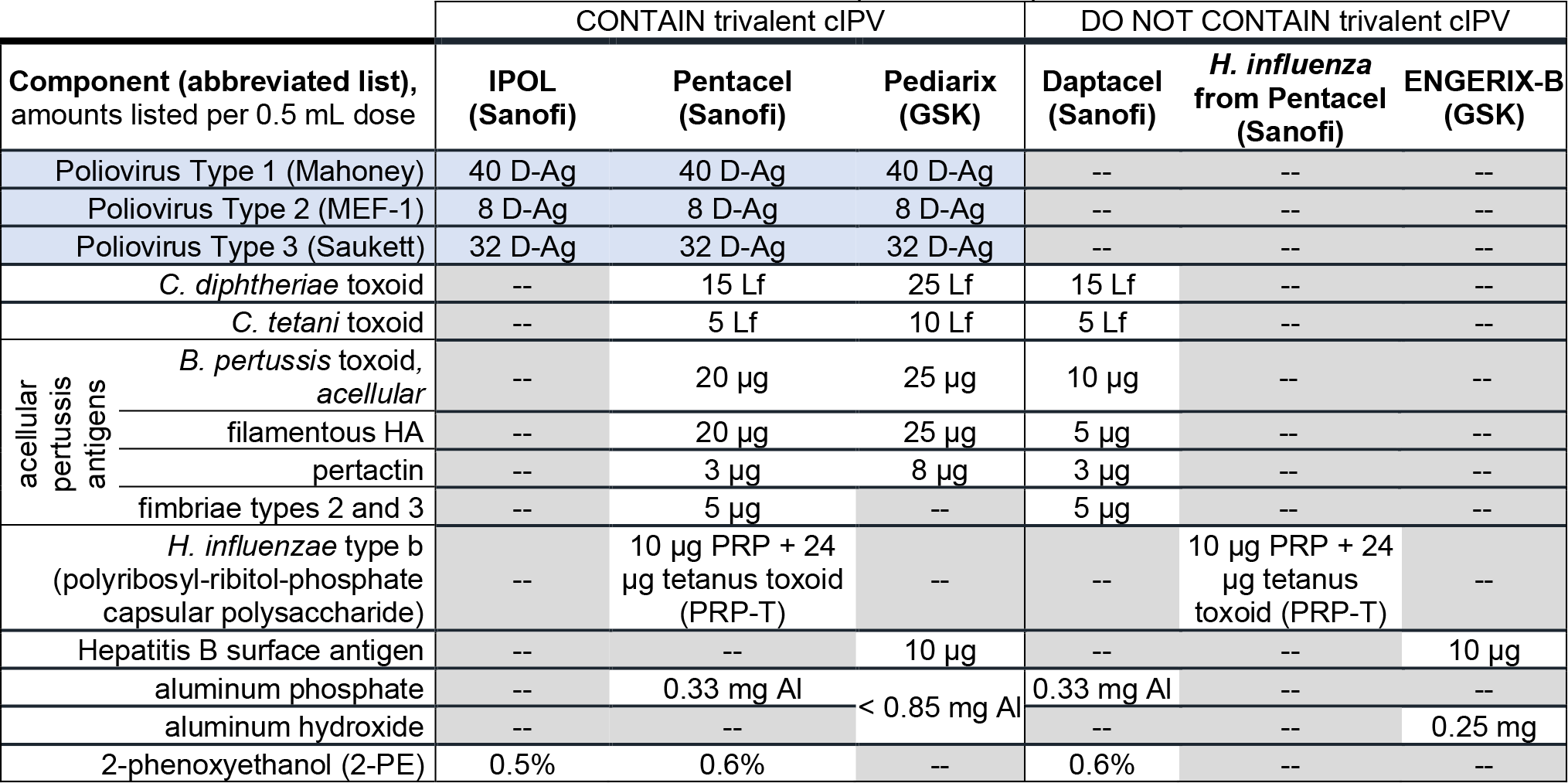
Components Present in Vaccines Used During Interference and Cross-Reactivity Testing, Excerpted from Relevant Vaccine Product Inserts,^13–17^ where “--” indicates the component is not present

The vaccines above were reconstituted as directed in the product insert if required, and diluted 4x in PBB prior to analysis (n=4 each). 2-phenoxyethanol was diluted to 0.6% v/v in PBB prior to analysis (n=3). Citrate buffer was prepared at 10% (w/v) and diluted 4x in PBB prior to testing (n=1). Imject Alum was diluted to 0.75 mg Alum/mL in PBB 2.0 or PBS (2 replicates each, n=4 total) prior to analysis. Cell culture matrix was evaluated using Vero cells prepared as described above. Cells were spun down, and the supernatant pulled off and diluted by 30% in PBB prior to analysis (n=1).

#### Vero Cell Culture Matrix

To mimic crude in-process samples relevant to vaccine bioprocessing, uninfected Vero cell culture was subjected to 3 freeze/thaw cycles and then clarified by centrifugation at 1000 x g for 5 min. The supernatant was removed without disturbing the resulting pellet. Eight (8) replicates of contrived trivalent sIPV were prepared in 50% Vero cell culture supernatant at 5.0/2.5/5.0 D-Ag units/mL for types 1/2/3, along with 8-point standard curves of the same trivalent sIPV sample prepared in either PBB or in the presence of 50% Vero cell culture. Vero cell medium without poliovirus was diluted with PBB and run as a negative control. The measured concentrations in the 8 replicates were then determined using each of the two standard curves, and accuracy and precision assessed against their respective calibrants. Accuracy was assessed as % of expected, and precision was assessed as % RSD of the replicates.

### 2.10 Trivalent cIPV-Containing Vaccines

IPOL, Pentacel, and Pediarix were tested and a list of critical ingredients in these vaccines excerpted from the vaccine product inserts can be found in **Table 1**.^13–17^ Vaccines and cIPV standard were diluted 4-fold in PBB, analyzed, and the % difference in the signals generated for the vaccine samples under investigation as compared to the WHO standard were calculated.

#### Citrate Desorption

Pediarix was mixed with 30% (w/v) sodium citrate tribasic dihydrate, pH=6 at 2:1 ratio and 3 identical aliquots placed in a 37°C incubator for 3 hours to desorb the polio antigens from the adjuvant. The treated samples were then centrifuged at 1500 x g for 5 min, and the supernatant was removed for analysis. The concentrations of each serotype in the supernatant were then measured against the cIPV WHO international standard. These values were compared to those obtained upon analysis of 3 aliquots of a Pediarix sample prepared in an inert matrix (no citrate) at the same nominal concentrations as the desorbed samples. The “no desorption” control samples were not spun down prior to analysis.

## 3. RESULTS AND DISCUSSION

### 3.1 VaxArray Poliovirus Assay Enables Serotype-Specific Detection of Types 1, 2, and 3

Monovalent sIPV materials were utilized to test VaxArray Polio Assay serotype specificity, as shown in Figure 2.

**Figure 2.**
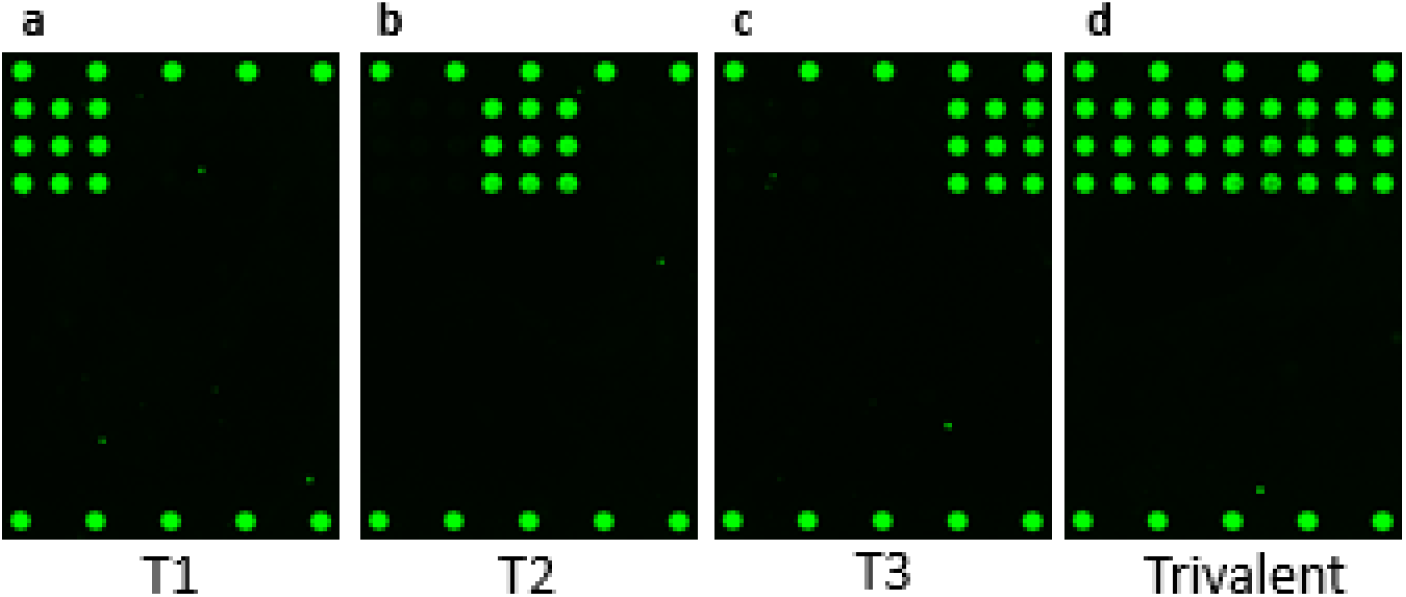
Example fluorescence microarray images demonstrating reactivity and specificity of the VaxArray Polio Assay after the incubation of monovalent sIPV samples in (a) T(1), (b) T(2), (c) T(3), and a trivalent mixture containing T(1), T(2), and T(3) in (d).

**Figures 2a–c** show representative fluorescence images of purified monovalent sIPV types 1, 2, and 3, respectively. These images qualitatively indicate that the serotype-specific antibodies generate strong signal responses to each of the D-antigen types and that the off-target antibodies do not produce signal above background. Quantitatively, the signal to background (S/B) ratios generated for types 1, 2, and 3 run monovalently were 50.4, 59.3, and 55.0, respectively, while all off-target antibodies in this analysis resulted in S/B ratios of < 1.1 indicating no appreciable positive signal. **Figure 2d** shows a representative fluorescence image when all 3 monovalent sIPV materials are mixed and tested as a trivalent mixture. With this specificity, the assay enables simultaneous analysis of all 3 serotypes in a trivalent mixture and provides a distinct advantage over an inherently singleplex standard plate-based ELISA.

To investigate the specificity of the assay for the D-antigen form of poliovirus, a forced degradation study was conducted. **Figure 3** shows the results of a thermal treatment of trivalent sIPV at 56°C as a function of time. The untreated material (0 minutes) produced typical VaxArray signals for the concentrations analyzed. As expected, after 15 minutes of thermal treatment the D-antigen form was thermally degraded, as reflected by greatly reduced signals approaching background signal (shown in grey), indicating specificity for the D-antigen form.

**Figure 3.**
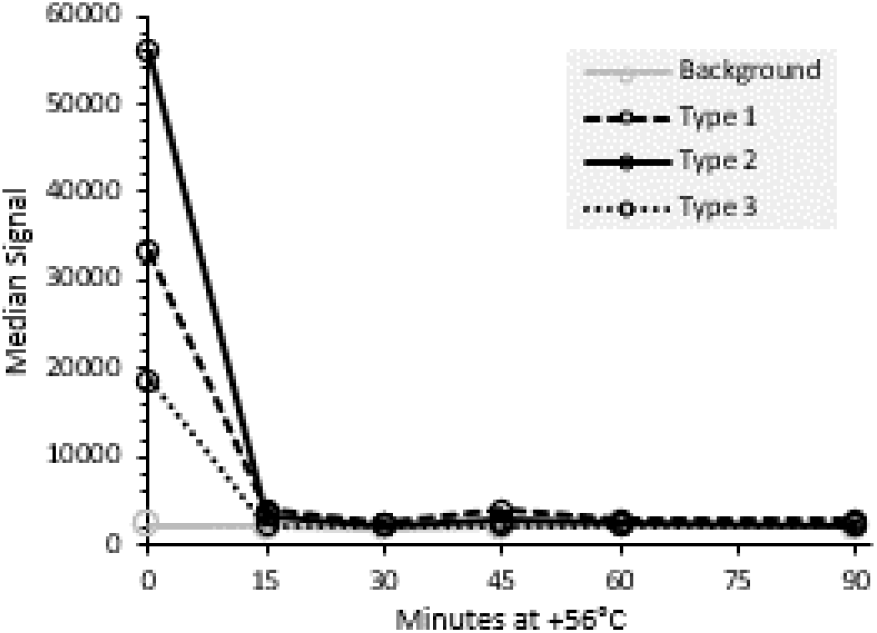
Thermal treatment of trivalent sIPV at 56ºC for varying amounts of time demonstrating assay specificity to the D-antigen form. For comparison to positive signals of T(1), T(2), and T(3), background signal (no antigen present) at each timepoint is shown as open grey circles with solid line.

### 3.2 VaxArray Poliovirus Assay is Quantitative with Sub-1 D-Antigen Unit/mL Sensitivity

**Figure 4** shows an 8-point dilution series for all 3 serotypes of a trivalent sIPV mixture. Eight replicates of three samples at concentrations near the expected LLOQ were analyzed against the standard curve (see the green, gray, and orange series in **Figure 4**). A similar analysis was also done for the monovalent sIPV materials to enable a comparison of metrics in the trivalent vs. monovalent formulations, and this complete analysis was also repeated for cIPV (IPOL vaccine). The LLOQ metrics determined for both sIPV and cIPV can be found in **Table 2**.

**Figure 4.**
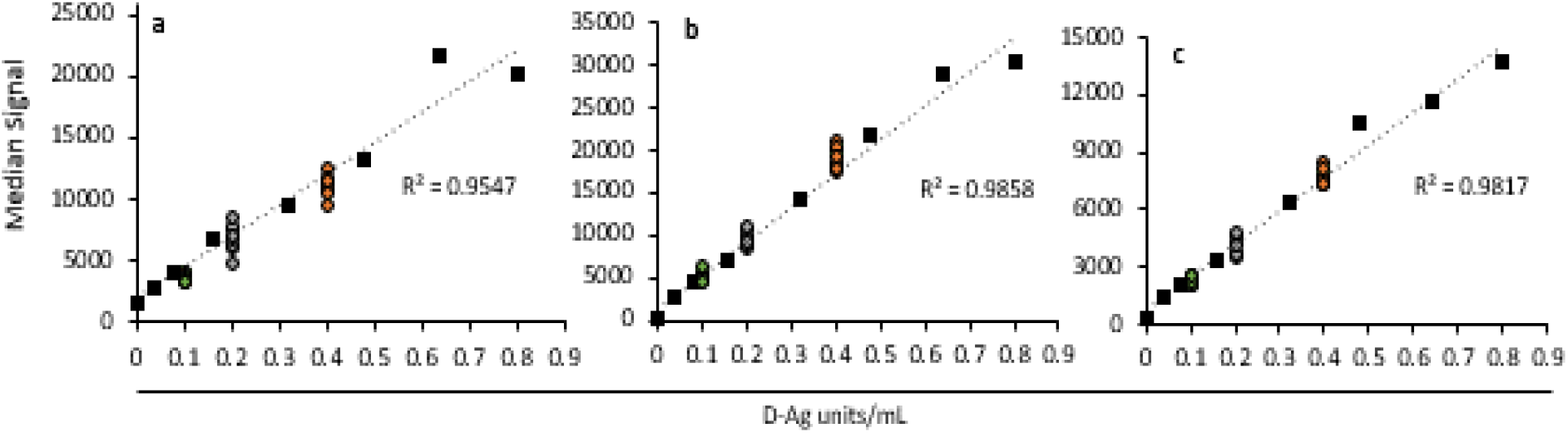
Response curves near the lower limit of quantification (LLOQ) for trivalent sIPV (black squares) for serotypes 1 (a), 2 (b), and 3 (c), along with 8 replicates analyzed at 0.1 D-Ag units/mL (green circles), 0.2 D-Ag units/mL (grey circles), and 0.4 D-Ag units/mL (orange circles). Y-axis is median fluorescence signal generated, and linear fits are dotted lines with the associated correlation coefficients (R^2^) indicated.

**Table 2:**
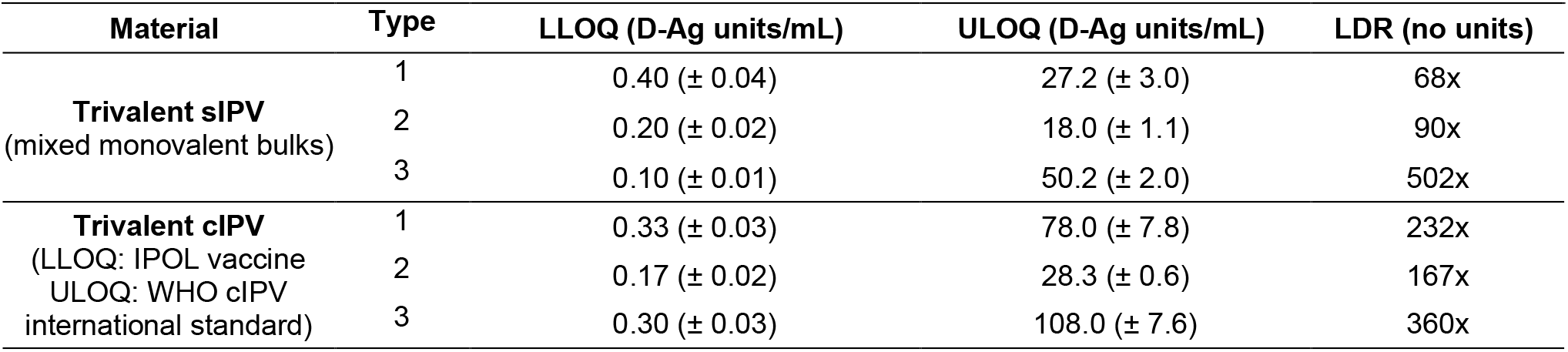
Analytical Sensitivity Metrics for Trivalent sIPV and cIPV

| Material | Type | LLOQ (D-Ag units/mL) | ULOQ (D-Ag units/mL) | LDR (no units) |
| --- | --- | --- | --- | --- |
| **Trivalent sIPV**<br>(mixed monovalent bulks) | 1 | 0.40 ($\pm 0.04$) | 27.2 ($\pm 3.0$) | 68x |
| | 2 | 0.20 ($\pm 0.02$) | 18.0 ($\pm 1.1$) | 90x |
| | 3 | 0.10 ($\pm 0.01$) | 50.2 ($\pm 2.0$) | 502x |
| **Trivalent cIPV**<br>(LLOQ: IPOL vaccine<br>ULOQ: WHO cIPV<br>international standard) | 1 | 0.33 ($\pm 0.03$) | 78.0 ($\pm 7.8$) | 232x |
| | 2 | 0.17 ($\pm 0.02$) | 28.3 ($\pm 0.6$) | 167x |
| | 3 | 0.30 ($\pm 0.03$) | 108.0 ($\pm 7.6$) | 360x |

Lower limits of quantification ranged from 0.17 to 0.40 D-Ag/mL, and the ULOQ ranged from ~18 to 108 D-Ag/mL. The linear dynamic range (LDR) was then defined as the ULOQ/LLOQ, with all values shown in **Table 2**.

Given that cIPV concentrations in final trivalent vaccines are typically 40/8/32 D-Ag per 0.5 mL dose for types 1/2/3 respectively, and that sIPV vaccines contain > 1.5 D-Ag per 0.5 mL dose in each serotype, the LLOQ is more than adequate for quantifying D-Ag content in final vaccine formulations. This excellent sensitivity is helpful when analyzing samples in a variety of crude matrices, as an upfront dilution minimizes any potential interferents. Furthermore, higher concentration samples such as those encountered in bioprocessing steps can easily be analyzed by diluting the sample to within the linear range as needed. We additionally examined VaxArray Polio Assay response curves for the OPV WHO international standards to confirm reactivity for applicability to this vaccine type (see Supplemental Information for details). While live-attenuated OPV is typically assessed via an infectivity measurement, there may be value in analyzing OPV for antigen assessment during bioprocess development and optimization given the rapid turnaround time.

### 3.4 Assay Exhibits Good Accuracy and Precision

Trivalent sIPV material at 5 D-Ag/mL in each serotype was used to evaluate user to user and day to day accuracy and precision. Three users performed the assay on 8 replicates alongside a standard curve of the same material. Each user repeated this analysis on 3 separate days (n=72 datapoints for each serotype in total). **Table 3** contains the accuracy and precision data generated, separated by serotype and user as well as combined over all 3 users. Accuracy values, expressed as % of expected or % recovery, ranged from 91% to 114%, with an overall average accuracy of 100 (± 8)%. This is well within a typical acceptable 80-120% recovery range. Precision values are expressed as the %RSD of the replicates in **Table 3** and ranged from 9% to 15%, with an overall average precision of 12 (± 2)%.

**Table 3:**
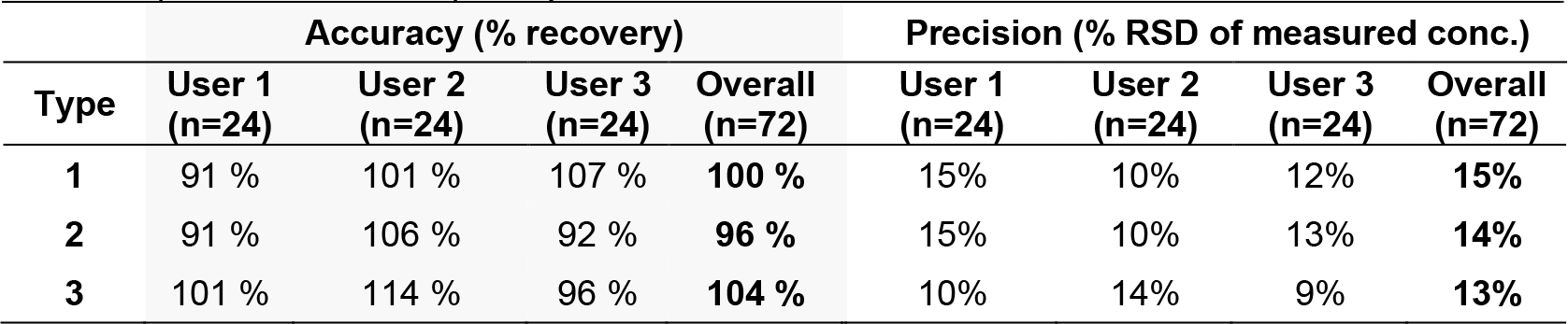
Accuracy and precision of quantitation of trivalent sIPV (5 D-Ag units/mL each serotype) over multiple users and multiple days

To compare assay accuracy and precision between sIPV and cIPV, samples of both were tested by a single user over a range of concentrations and over multiple assay setups, as shown in **Table 4**. For sIPV, 8 replicate samples at 1.5, 5.0, and 10.0 D-Ag units/mL were tested in each of 3 separate assay setups. For cIPV, a similar analysis was conducted using 1.5/0.38/1.35, 5/1.27/4.51, and 10/2.55/9.02 D-Ag units/mL for the low, medium, and high concentrations of types 1/2/3, respectively.

**Table 4:** Accuracy and precision of quantitation of trivalent sIPV (1.5, 5.0, and 10.0 D-Ag units/mL) and IPOL vaccine (trivalent cIPV, 1.5/ 0.38/ 1.35, 5/ 1.27/ 4.51, and 10/ 2.55/ 9.02 D-Ag units/mL) at low, medium, and high concentrations, n=24 for each measurement (single user, 8 replicates of each sample run in each of 3 separate assay setups)

|  |  | Accuracy (% recovery) |  |  | Precision (% RSD of measured conc.) |  |  |
| --- | --- | --- | --- | --- | --- | --- | --- |
|  | Type | Low | Med | High | Low | Med | High |
| sIPV | 1 | 105% | 99% | 96% | 17% | 11% | 13% |
|  | 2 | 97% | 106% | 95% | 13% | 10% | 12% |
|  | 3 | 92% | 114% | 88% | 14% | 11% | 11% |
| cIPV | 1 | 114% | 108% | 108% | 16% | 6% | 9% |
|  | 2 | 114% | 113% | 110% | 12% | 13% | 10% |
|  | 3 | 115% | 106% | 92% | 20% | 14% | 16% |

Accuracy values for sIPV, ranged from 88-114% recovery, with an average of 99 (± 8)%. Accuracy for cIPV ranged from 92 to 115%, with an average of 109 ± 7%, and there were no notable differences as a function of concentration. Together the data indicate similar performance across the linear dynamic range for both cIPV and sIPV. Precision values, as shown in **Table 4**, were also similar between sIPV and cIPV. Expectedly, slightly higher % RSD was observed at the lowest concentration in many cases for both sIPV and cIPV. All % RSD values were at or below 20%, with an overall average precision of 12 (± 2) % RSD for sIPV and 13 (± 4) % for cIPV, indicating reasonable precision of replicate measurements. Of note, the overall precision was similar to those for the multi-user, multi-day sIPV study summarized in **Table 3**.

### 3.5 Assay with ≤ 3-Hour Time to Result has Improved Accuracy over 3-Day ELISA

A trivalent mixture of monovalent bulk sIPV analyzed at a variety of dilutions alongside a trivalent WHO sIPV standard curve was utilized to compare accuracy and precision of the VaxArray Polio Assay to the 3-day plate-based ELISA described by Kouiavskaia et al. that uses the same sample capture and label antibodies.^12^ **Figure 5** shows the % recovery (accuracy) and % RSD (precision) of replicate measurements for both VaxArray and ELISA on samples spanning concentration ranges from ~20 D-Ag units/mL to 0.1 D-Ag units/mL. **Figures 5a, b, and c**, show T(1), T(2), and T(3), respectively (n=6 for each concentration for each method). The dotted line represents 100% recovery, and the light shaded box highlights measurements falling within 85-115% of the expected result. As seen in **Figure 5**, for all 3 serotypes, 5 of the 7 (~71%) VaxArray average measurements are within 15% of the expected result; whereas only 2 of the 7 ELISA (~28%) average measurements are within 85-115% of expected for T(1) and T(2), and only a single sample is within these bounds for T(3), indicating a higher accuracy for the VaxArray Polio Assay. The combined accuracy over all the samples tested is shown in the solid and hashed blue bars at the right of **Figure 5**, with the overall VaxArray values producing 110%, 100%, and 100% average recovery for types 1, 2, and 3, respectively. In contrast, the ELISA resulted in 84%, 96%, and 78% average recovery for types 1, 2, and 3.

**Figure 5.**
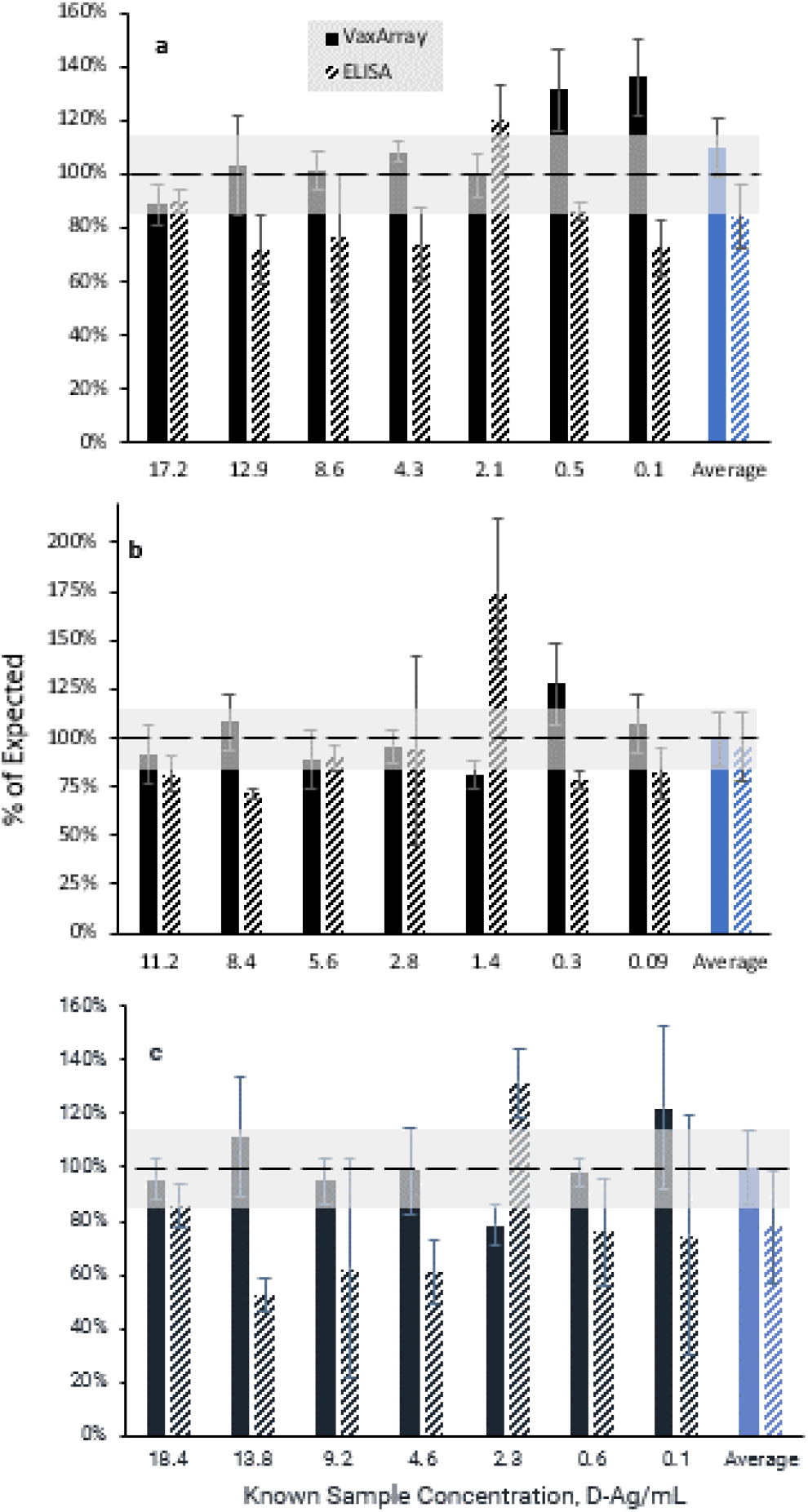
Comparison of % expected concentration between VaxArray Polio Assay and plate-based ELISA. D-Ag concentrations are grouped as shown on the x-axis, with the overall average % recovery over all concentrations investigated shown in the blue bars at right. (a) T(1), (b) T(2), and (c) T(3), with VaxArray Polio Assay measurements shown as solid black bars, and ELISA measurements shown as hashed black and white bars. Error bars represent % RSD of 3 replicate measurements over 2 separate experiments (n=6 for each method at each concentration).

In terms of precision, the % RSD of the replicate measurements are represented by the error bars in **Figure 5**. VaxArray generated average % RSDs of 11%, 14%, and 15% for T(1), T(2), and T(3), whereas ELISA produced average % RSD of 12%, 17%, and 21% for T(1), T(2), and T(3). These precision values indicate similar precision was achieved by both methods in this experiment. A direct correlation between the ELISA and VaxArray data resulted in correlation coefficients (R^2^) for types 1, 2, and 3 of 0.93, 0.92, and 0.84, respectively (data not shown). Importantly, the plate-based ELISA has a 3-day time to result and is only capable of analyzing a single serotype per well, such that separate wells must be coated for each serotype under analysis. In contrast, the VaxArray Polio Assay’s multiplexing capability allows for simultaneous quantification of all 3 serotypes in a single assay, and results were obtained same-day (3 hours including sample preparation). We also note that the VaxArray time to result can be further shortened to less than 1 hour using the alternative ArrayMax orbital shaker, (see Supplemental Information for details), further enhancing the utility of the assay.

### 3.6 Assay is Suitable for Bioprocess Samples and Multivalent Drug Product

#### 3.6.1 Quantification in Crude Cell Culture Matrix

To demonstrate applicability to crude in-process samples relevant to vaccine bioprocessing, trivalent sIPV was added to exhausted, clarified Vero cell culture medium and analyzed. **Table 5** shows the accuracy and precision in the back-calculated concentration achieved for 8 replicate analyses of mock “crude” trivalent sIPV. The same trivalent sIPV mixture was used as a calibrant within a standard curve prepared in a clean matrix (PBB) or prepared in the same 50% Vero cell culture matrix as the replicates. Observed precision of the replicates ranged from 6 to 11 % RSD and was similar when comparing the “matrix matched” vs. non-matrix matched calibration approach. In addition, precision values were similar to those obtained for both sIPV and cIPV, as shown in **Tables 3** and **4**. When calibrating against the clean/non-matrix matched standards (left columns of **Table 5**), % recovery was 90% and 86% of expected for T(1) and T(3), respectively. However, % recovery for T(2) was quite low at only 64% of expected. In contrast, when calibrating against the matrix-matched standard curve (righthand columns of **Table 5**), % recovery for all 3 types was ≥ 84% of the expected result, indicating that utilizing a matrix-matched standard curve can significantly improve quantitation. These data indicate that the assay should be suitable for the rapid analysis of crude in-process samples provided that a matched calibrant is included for accurate quantitation.

**Table 5:** Quantification of trivalent sIPV in 50% Vero cell culture supernatant against calibration curve of trivalent sIPV in either PBB or 50% Vero cell culture supernatant (n=8 replicates)

| Type | NOT Matrix Matched<br>calibrated against trivalent sIPV in PBB |  | Matrix Matched<br>calibrated against trivalent sIPV in Vero cell matrix |  |
| --- | --- | --- | --- | --- |
|  | Precision (% RSD) | Accuracy (% recovery) | Precision (% RSD) | Accuracy (% recovery) |
| 1 | 8% | 90% | 6% | 86% |
| 2 | 6% | 64% | 6% | 84% |
| 3 | 11% | 86% | 10% | 87% |

#### 3.6.2 Potential Interferents Do Not Produce False Positive Signal

A series of studies was conducted to evaluate the effect of common vaccine additives on the assay. Specifically, Daptacel, EngerixB, and ActHIB (the *H. influenza* component present in Pentacel) vaccines were tested in the absence of polio virus (see **Table 1** for a list of components). The assay does not result in appreciable false positive signal in the presence of these substances, with S/B ratios less than 1.5 for all substances tested (data not shown). Additives such as 2-phenoxyethanol and alum adjuvant also yield no appreciable assay signal above background. Lastly, exhausted, clarified Vero cell culture supernatant (relevant for crude, in-process samples as discussed previously) and citrate buffer (relevant for use of citrate-based antigen desorption protocols) both analyzed in the *absence* of poliovirus did not produce any appreciable signal, as indicated by signal to background ratios ≤ 1.5.

#### 3.6.3 Analysis of cIPV in Combination Vaccines

To assess assay quantification of poliovirus D-antigen in final trivalent cIPV-containing poliovirus vaccines and combination vaccines, three vaccines were tested (see **Table 1).** All 3 vaccines were assumed to contain the stated minimum concentrations of 40/8/32 D-Ag units per dose for polio types 1/2/3 (exact concentrations unknown) as they were within their stated shelf life and were therefore compared to the same nominal concentrations in the WHO cIPV international standard. **Figure 6** compares the % expected signals generated for T(1), T(2), and T(3) for the IPOL and Pentacel vaccines compared to the WHO cIPV international standard.

**Figure 6.**
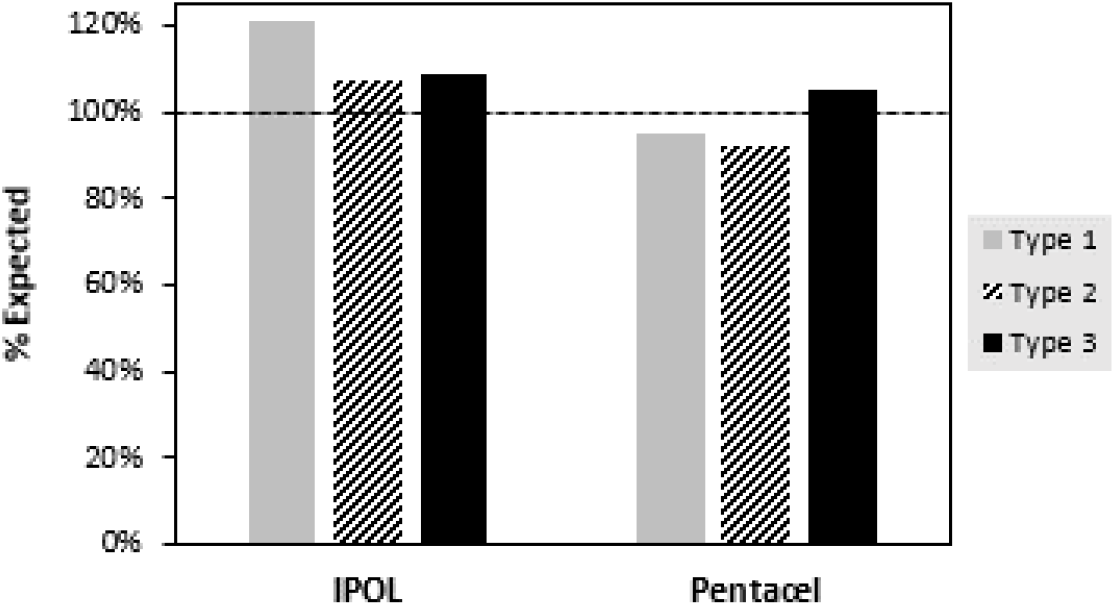
VaxArray Polio Assay measurements in trivalent cIPV-containing vaccines. IPOL and Pentacel (see **Table 1** for list of components in each; both contain a minimum 40/8/32 D-Ag units and were analyzed by VaxArray Polio Assay and compared to the same nominal concentrations of cIPV WHO international standard, with results shown as of expected concentration. T(1) results are shown as solid grey bars, T(2) is hashed black and white bars, and T(3) is solid black bars as shown.

For IPOL, all 3 types produced between107 and 120% of the expected concentration, and Pentacel produced % differences from 92 to 105% of the expected concentration depending on type. Given that this comparison represents the *minimum* concentration present in the vaccines, % values greater than expected may represent a slight overfill of the vaccine, which is a common practice. Therefore, the assay was considered to produce reasonably accurate measurements for both IPOL and Pentacel as compared to the WHO cIPV standard. Based on an analysis of the components present in IPOL and Pentacel (see **Table 1)**, we noted that IPOL is not adjuvanted, and Pentacel contains 0.33 mg Al as aluminum phosphate. Analysis was also conducted of the Pediarix vaccine which contains < 0.85 mg/mL of Al as a mixture of aluminum phosphate and aluminum hydroxide. This analysis (see righthand bars in **Figure 7**) resulted in a much lower % of the expected concentration, with average % expected of 58, 64, and 67% for T(1), T(2), and T(3), respectively. This may indicate an interference from the aluminum hydroxide component, given that Pentacel containing aluminum phosphate showed good accuracy. Vaccine antigens are often adsorbed to adjuvants to enhance the immune response, with aluminum-containing adjuvants being most commonly utilized.^18,19^ It is well-known that a desorption step prior to analysis by most *in vitro* assays, including ELISAs, is required due to the potential for interference.^20,21^ A variety of desorption methods can be used, including manipulation of pH,^22^ addition of surfactants,^23,24^ and addition of citrate buffer to dissolve the adjuvant.^18,19,25,26^

**Figure 7.**
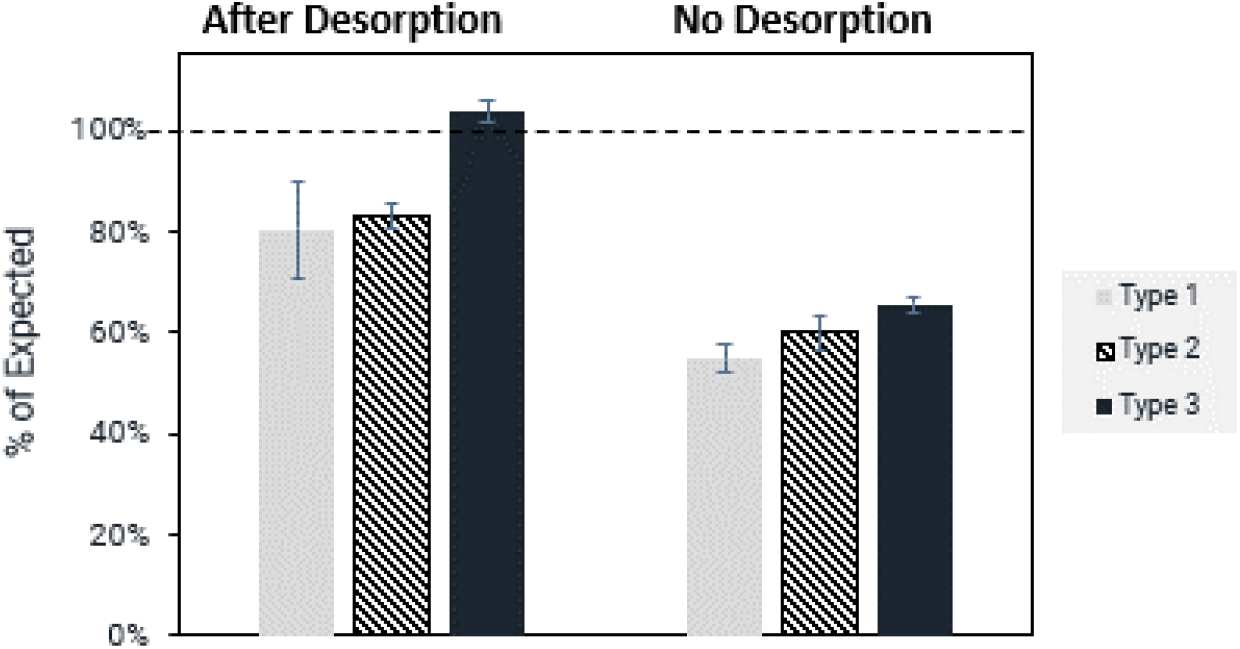
VaxArray Polio Assay measurements of Pediarix trivalent cIPV-containing vaccine indicating significantly improved recovery after citrate desorption. Results are shown as % of expected concentration based on comparison to the same nominal concentrations of cIPV WHO international standard, with error bars shown as the % RSD of triplicate measurements. T(1) results are shown as solid grey bars, T(2) is hashed black and white bars, and T(3) is solid black bars as shown.

To determine if a citrate desorption step would allow for better recovery and accuracy for Pediarix by the VaxArray Polio Assay, we subjected the Pediarix vaccine to 37°C for 3 hr in the presence of citrate buffer at final nominal concentrations of 14/2.8/11.2 D-Ag/mL for types 1/2/3 and compared the assay response to Pediarix prepared at the same concentrations but *not* subjected to desorption (no citrate present). **Figure 7** shows comparative responses of the desorbed and untreated samples. The untreated aliquots (no desorption) shown in **Figure 7** produce measured concentrations of only 58 to 67% of expected, as noted previously. After citrate desorption, analysis of the supernatant indicates that all 3 serotypes produce average concentrations within ± 20% of expected. While this citrate protocol could certainly be further optimized, or alternative methods of desorption investigated, these data indicate that the VaxArray Polio Assay can be utilized on samples that have undergone a citrate desorption step to improve recovery in adjuvanted samples.

## 4. CONCLUSION

Development and validation of standardized analytical tools that can shorten the time to result for characterization of vaccines is an important step forward and can significantly enhance operational efficiency. While standard ELISAs are commonly utilized to measure D-antigen content of inactivated poliovirus vaccines and efforts are underway to improve and standardize these existing assays, ELISAs typically have long assay times of 1 to 3 days, suffer from lab-to-lab variability, are not amenable to multiplexing all antigens in a multivalent vaccine in a single test, and utilize significant amounts of reagents compared to microscale approaches. In contrast, the VaxArray Polio Assay has been developed to use the same serotype-specific capture antibodies and universal label antibody recently validated in a standard ELISA platform, but offers a simple, rapid, high throughput method for IPV and combination vaccines that offers the benefits of standardized reagent kits, multiplexing, use of 10-100x less capture reagent than ELISA, and a rapid time to result. The work presented herein demonstrates the VaxArray Polio Assay achieves similar or improved analytical performance relative to the polio D-antigen ELISA but has a ~25x faster time to result. Importantly, the VaxArray assay works well for combination vaccines that contain polio D-antigen and it is compatible with common vaccine additives, adjuvants, and crude matrices applicable to bioprocess samples. We hope this new tool will be utilized by poliovirus vaccine manufacturers worldwide and validated in their labs for characterization and release testing as part of the critical effort to update and adapt poliovirus vaccine manufacturing for increased safety in a post-eradication world.

## ACKNOWLEDGEMENTS

This work was supported, in whole or in part, by the Bill & Melinda Gates Foundation [INV-004629]. Under the grant conditions of the Foundation, a Creative Commons Attribution 4.0 Generic License has already been assigned to the Author Accepted Manuscript version that might arise from this submission. PATH provided monovalent Sabin IPV materials (serotypes 1, 2, and 3) from two different unnamed manufacturers, as well as research quantities of the four human anti-D-antigen polio monoclonal antibodies used for serotype-specific capture and detection labeling to InDevR under a formal research collaboration agreement. We also acknowledge Raul Hernandez at Global Sourcing Initiative for sourcing of the on-market vaccines utilized herein.

## AUTHOR CONTRIBUTIONS

Erica Dawson: project administration, supervision, methodology, resources, formal analysis, writing-original draft, visualization; Kathy Rowlen: conceptualization, project administration, resources, writing-review and editing; James Johnson: methodology, investigation, validation, formal analysis, writing-review and editing; Tianjing Hu: methodology, investigation, validation, formal analysis, writing-review and editing; Amber Taylor: supervision, methodology, formal analysis, visualization, writing-review and editing; Caitlin McCormick: investigation, validation, formal analysis, writing-review and editing; Keely Thomas: investigation, validation, formal analysis, writing-review and editing; Rachel Gao: investigation, validation, formal analysis, writing-review and editing

## SUPPLEMENTAL INFORMATION

### Oral Poliovirus Reactivity

Oral Poliovirus Vaccine (OPV) WHO international standards for OPV1, 2, and 3 were obtained from NIBSC (16/196, 15/296, and 16/202) with stock concentrations ranging from 10^6^-10^7^ TCID_50_/mL. Based on literature, the concentrations in terms of D-Ag/mL are expected to be from ~0.3 to 3 D-Ag/mL.^1,2^ A 10-fold dilution series for each serotype was analyzed monovalently (n=1 each dilution due to limited sample), with the highest concentration samples not diluted in PBB to enable testing at maximum concentration. A 6-hr antigen incubation time was utilized to ensure adequate signal from these low concentration samples. A linear fit was applied to each response curve to assess linearity, with the error bars expressed as ± 1 standard deviation of the median fluorescence signal. For OPV1, the highest concentration standard was omitted from the linear fit due to fluorescence saturation. We note that InDevR is a polio essential facility (PEF) in accordance with the US National Authority for Containment of Poliovirus (NAC) housed at the US Centers for Disease Control and Prevention (Atlanta, GA).

**Figure S1** shows the median signals generated by each serotype. Due to limited antigen material, a single replicate was run for each concentration with error bars representing ± 1 standard deviation of the 9 replicate microarray spots. In **Figure S1a**, the highest concentration sample for serotype 1 was fluorescently saturated and therefore not included in the fit. All three serotypes demonstrated a robust linear response curve, with R^2^ values of 0.95, 0.99, and 0.94 for T(1), T(2), and T(3), respectively. These results demonstrate that the assay can be applied to quantify OPV samples of sufficient concentration with the appropriate standards. Given that infectivity, such as TCID_50_, is required as a release assay for this live-attenuated vaccine, it is unlikely that the VaxArray Polio Assay will be a suitable release assay for OPV, but such a rapid method has utility during bioprocess development and optimization to provide complementary, actionable information with a same day turnaround time.

**Figure S1.**
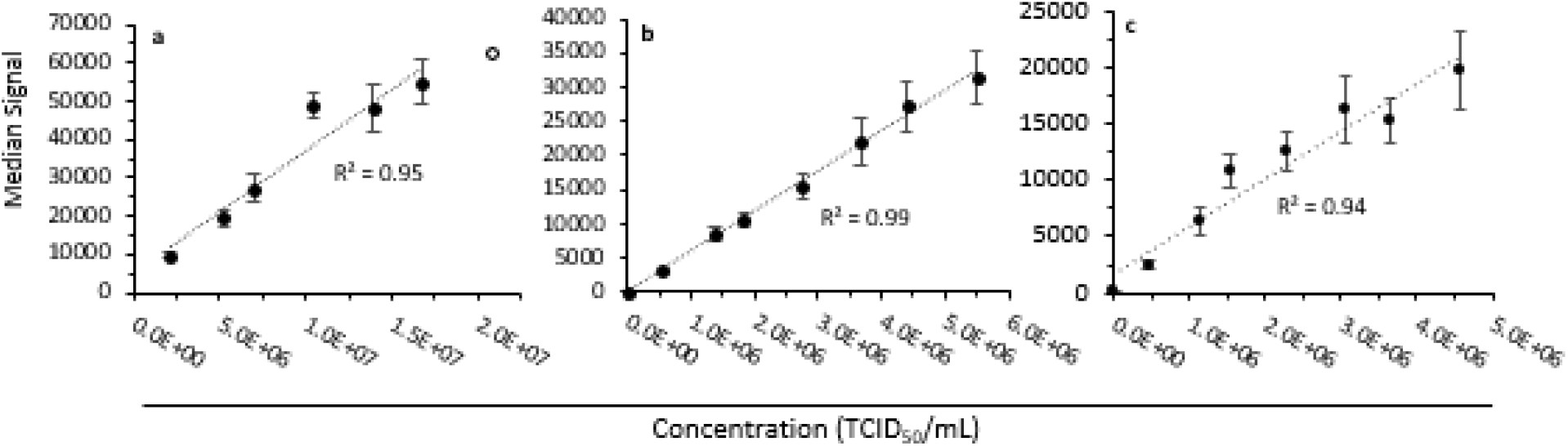
Monovalent OPV response curves for (a) T(1), (b) T(2), and (c) T(3). Y-axis is median fluorescence signal generated, and linear fits to the response curves for each serotype are shown as dotted lines with the associated correlation coefficients (R^2^) indicated. In (a), the highest concentration point is omitted from the fit due to fluorescence saturation. Because a single replicate of each sample was analyzed, error bars represent ± 1 standard deviation of the 9 replicate microarray spots for each capture.

### sIPV Time to Result Reduction Studies

To further minimize the assay time to result, additional testing was performed using the ArrayMax orbital shaker (VX-6212, InDevR, Inc.) instead of a standard orbital shaker during antigen incubation and detection labeling. Monovalent sIPV materials were combined to 100/50/100 D-Ag/mL starting concentrations for types 1/2/3 and used to create a standard curve of 15 dilutions down to 0.03 D-Ag/mL which was analyzed in triplicate. Slides were placed on the ArrayMax platform, 45 μL of samples added to the microarrays, and shaken at 700 rpm for 15 min. Samples were then removed, and 45 μL of detection label was added and allowed to shake at 700 rpm for an additional 15 min. Slides were then processed and imaged as previously described.

While the VaxArray Polio Assay was initially developed for use with a standard large orbit (20 mm diameter) shaker (such as SCI-O180-S, Scilogex) with a 3-hour time to result, here we demonstrate that it is quite feasible to obtain essentially equivalent results in under 1 hour for the VaxArray Polio Assay using a small orbit (1mm) shaker (ArrayMax, InDevR, Inc.) to improve mass transport. For the assay performed with the large-orbit shaker, 2 hr antigen incubation and 30 min labeling steps were used (2.5 hours overall on shaker). For the assay performed using the small-orbit shaker, 15 min antigen incubation and 15 min labeling steps were used (30 minutes on shaker). **Figure S2** shows comparative response curves for both the large-orbit shaker (solid lines), and the small-orbit ArrayMax shaker (dotted lines), with the smoothed fits intended only to guide the eye.

**Figure S2.**
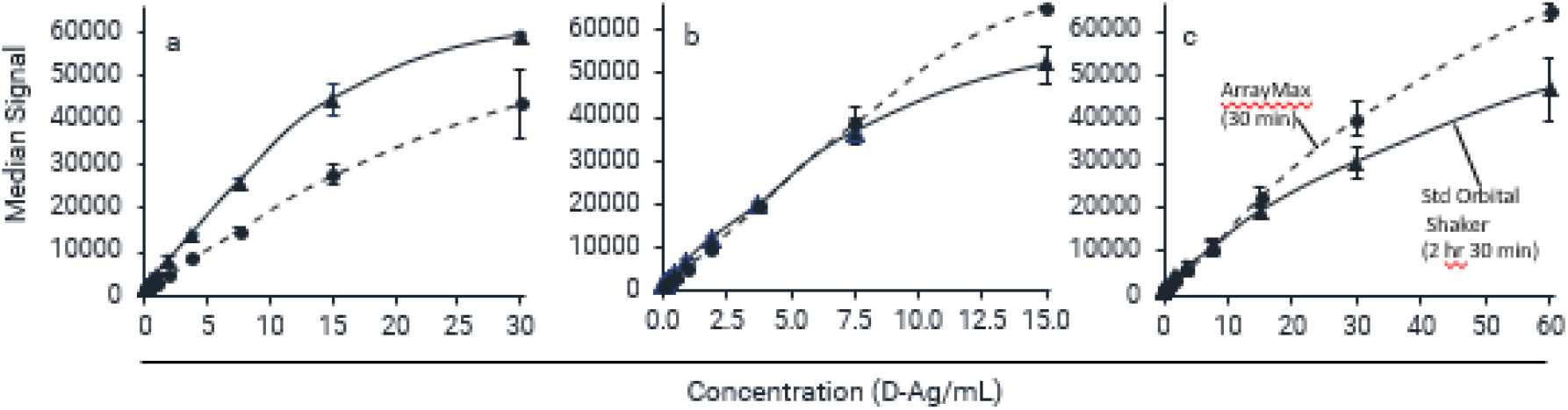
Comparison of response curves using standard orbital shaker and ArrayMax orbital shaker to improve mass transport and reduce assay time to result. T(1), T(2), and T(3) are shown in panels (a), (b), and (c), respectively. Standard orbital shaker with 2-hour antigen incubation and 30-min detection labeling times is shown as filled triangles with a solid black line to guide the eye, and ArrayMax orbital shaker with 15-minute antigen incubation and 15-min detection labeling times is shown as the filled circles with the dotted line to guide the eye. Error bars shown are ± 1 standard deviation of 3 measurements. Both datasets were collected using the same 100 ms exposure time.

A similar signal response (either slightly lower or higher than the standard assay) was generated for all 3 serotypes while reducing the overall incubation time by 5x, and the same exposure time was utilized during fluorescence imaging for both methods. In addition, the error bars shown in **Figure S2** (± 1 standard deviation of the mean of 3 independent experiments) illustrate that similar precision is obtained for both methods. Achieving a rapid time to result is of particular importance for bioprocess samples in which a rapid turnaround time is desirable to produce actionable information that can inform bioprocess improvements or raise alerts during bioprocess monitoring. While the analyses conducted here were on purified poliovirus monovalent bulk vaccine materials, we have demonstrated for other VaxArray product lines (data not shown) that ArrayMax can be utilized to reduce the time to result for crude samples as well with similar results.

